# Embolization-on-a-chip: Novel Vascularized Liver Tumor Model for Evaluation of Cellular and Cytokine Response to Embolic Agents

**DOI:** 10.1101/2025.03.30.646225

**Authors:** Huu Tuan Nguyen, Zuzana Tirpakova, Arne Peirsman, Surjendu Maity, Natashya Falcone, Satoru Kawakita, Danial Khorsandi, Ahmad Rashad, Neda Farhadi, Kalpana Mandal, Menekse Ermis, Rondinelli Donizetti Herculano, Alireza Hassani Najafabadi, Mehmet Remzi Dokmeci, Natan Roberto De Barros, Ali Khademhosseini, Vadim Jucaud

## Abstract

**Background:** Embolization is a well-established treatment modality for liver cancer. However, traditional embolization agents are limited by inefficient delivery and aggregation in blood vessels. Novel shear-thinning hydrogels (STH) have been developed to address the need for safer and more effective local delivery of embolic agents and therapeutics.

**Objective:** We aim to evaluate the efficacy of novel embolic agents such as STH using a human-relevant in vitro model that recapitulates human hepatocellular carcinoma capillary networks.

**Methods:** A vascularized human liver-tumor-on-a-chip model was developed to assess embolic agent performance. The effects of drug-eluting STH (DESTH) on tumor cell viability, surface marker expression, vasculature morphology, and cytokine responses were evaluated. To study the effects of embolization on microvasculature morphology independent of the chemotherapy compound, we assessed the effect of different drug-free embolic agents on the vascular tumor microenvironment under flow conditions.

**Results:** DESTH treatment induced tumor cell death, downregulated the expression of Epithelial Cell Adhesion Molecules (EpCAM) in HepG2, increased levels of cytokines such as Interleukin-4 (IL-4), Granulocyte-macrophage colony-stimulating factor (GM-CSF), and Vascular Endothelial Growth Factor (VEGF), and decreased albumin secretion. Furthermore, different embolic agents exert distinct effects on microvascular morphology, with STH causing complete regression of the microvascular networks.

**Conclusion:** This vascularized liver tumor-on-a-chip model enables human-relevant, real-time assessment of embolic agent efficacy and vascular response, paving the way for the development of innovative and effective embolization therapies for liver cancer.

## INTRODUCTION

Hepatocellular carcinoma (HCC) accounts for approximately 90% of all liver cancer cases and is projected to cause more than a million new cancer cases by 2030 (1). Although the liver receives blood from two inflows: the portal vein and the hepatic artery, embolization involves the deliberate occlusion of arteries and capillaries with embolic agents and chemotherapeutic drugs, disrupting the blood supply to the target tumor tissue (2, 3). Therefore, transcatheter arterial chemoembolization (TACE) has been used as the first-line treatment for patients with intermediate stages of HCC (2, 4). TACE treatment locally releases chemotherapy agents such as doxorubicin (DOX), cisplatin, or mitomycin C in an iodinated lipid compound (5). Current efforts in developing new TACE compounds have focused on biodegradable, multifunctional nanomaterials and nanobeads (6). Shear-thinning hydrogels (STH) have been developed as injectable therapeutics because they can reduce viscosity under shear stress and recover when the stress is removed (7, 8). Their mechanical properties allow for seamless injection and delivery through needles and catheters. The drug-eluting STH (DESTH), in combination with immunotherapy agents previously developed by our group, has been used for several TACE applications (9).

Currently, animal models (i.e. rats and rabbits) are utilized to assess the efficacy and safety of embolic agents (10, 11). One challenge in animal testing is replicating the pathophysiology of human HCC and ensuring reproducibility at the molecular, cellular, and tissue levels. Also, animal models have blood vessel sizes and anatomy that are different from those of humans. Therefore, the effects of embolic agents in animal models are difficult to translate to human patients (12). Since animal models poorly represent human physiology, they are no longer mandatory for drug response evaluation by the United States Food and Drug Administration and the European Medicines Agency (13).

Another approach to TACE testing is using decellularized liver models (14, 15). These models enable visual and quantitative evaluation of embolic agents with retained native extracellular matrix and organ structures, but they do not recapitulate the cellular or cytokine responses to the treatments. Therefore, there is an urgent need for advanced *in vitro* models that more closely mimic the tumor microenvironment (TME) to evaluate the response of blood vessels and tumor cells to embolic agents.

*3D in vitro* human-based liver cancer models, such as spheroids, which are derived from cell lines, and organoids, which are generated from stem cells or patient tissues, allow the recreation of tumoral tissue structure and cell interactions (16). However, they cannot recapitulate fluid shear stress, hydrostatic pressure, tissue deformation, and perfusion of blood or nutrient-rich medium flowing through endothelium-lined vasculature (17, 18). Organ-on-a-chip (OOC) is an emerging platform implying engineered or natural miniature tissues grown in a controlled cell microenvironment while allowing perfusion flows into the tissue structures (19). Recently, researchers have advanced the development of various *in vitro* vascular beds for the studies of vascular biology, tumor microenvironments, and drug screening (20–22). The introduction of a vascular flow system on a chip has facilitated insights into the early and late phases of TACE treatment responses, including observations of drug release (23), physical parameters about drug elution (24) and spatial distribution, as well as the deformation of the spheres (25) under the fluidic conditions (26). Özkan et al. developed a complex vascularized HCC-OOC composed of HCC, endothelial, stellate, and Kupffer cells. The model consisted of one vascular channel simulated TACE in the channel seeded with vascular endothelium, allowing the monitoring of metabolic activity and changes in vascular permeability (27). Currently, no study has simulated TACE effects in liver models with capillary-like vessels.

In this study, we present a novel vascularized liver tumor model designed specifically to evaluate vessel remodeling and cell death in response to chemoembolic agents. Our model incorporates a tumor spheroid surrounded by perfusable capillary-mimicking microvascular networks in a microfluidic OOC system. We mimic the embolization by occluding the inflow of perfusable microvasculatures with embolic agents loaded via a catheter. We observed vessel regression caused by embolic agents and release of chemotherapy agents. The responses to embolization alone or with chemotherapy were assessed using cytokine release profile and cell surface marker staining. Finally, this model enabled the evaluation of various embolic agents, paving the way for their optimization and selection in future TACE therapies.

## MATERIALS AND METHODS

### Experimental design

The focus of the study was to create a functional vascularized device with an embedded liver cancer spheroid for *in vitro* chemoembolization testing. We adapted the design from a previously established integrative vascular bed (iVas) device to construct the vascularized tumor model (22). iVas consists of a fully perfusable microvascular bed surrounding an empty well that allows subsequent incorporation of grafted tissues. An HCC tumor spheroid was then placed into the well that was surrounded by the vascular bed connected to two media channels. A chemoembolization agent was introduced through one of the channels via a catheter to mimic the *in vivo* delivery of an embolization agent. Since the chemotherapy agent DOX is autofluorescent with a maximum excitation of 470 nm and an emission wavelength of 560 nm, it enabled the detection of its release throughout the device via fluorescence microscopy. The rate of chemotherapy release was compared with simulation test results. The cellular response of the vascularized tumor model was assessed using time-point live imaging and immunostaining. The cytokine response was characterized using ELISA and Luminex cytokine profiling. In the flow experiment, embolization agents, without chemotherapy agents, were used to assess the direct impact of embolization on the vasculature. The device was connected to a pump to generate a controlled media flow through the channels. Flow experiment data were collected four days after embolization. Vascular regression was quantified using MATLAB (MathWorks, USA) and ImageJ image processing software (28).

### Cell Culture

Human Umbilical Vein Endothelial Cells (HUVECs), either non-fluorescent (Lonza, USA) or GFP-labeled (Angioproteomie, USA), were cultured in EGM-2 Endothelial Cell Growth Medium supplemented with all the components in the BulletKit (Lonza, USA), except Heparin, which was used at one-quarter of the original concentration. The passage number of HUVECs did not exceed 7 (p7). Normal human lung fibroblasts (FBs, Lonza) were cultured in low-serum Fibroblast Growth Medium 2 (PromoCell). Lung fibroblasts promote HUVEC’s microvasculature formation *in vitro* through paracrine signaling (29). Mesenchymal stem cells (MSCs) were purchased from ATCC (Cat. No. PCS-500–012) and cultured in low glucose DMEM (Gibco, USA) containing 10% FBS. Hepatocellular carcinoma cells (HEPG-2, ATCC) were cultured in Eagle’s Minimum Essential Medium (EMEM) from Corning, USA, supplemented with 10% fetal bovine serum (FBS) (Gibco) and 1% Penicillin-Streptomycin (Gibco).

### Spheroid formation

Hepatocellular carcinoma cells HepG2 (ATCC) were seeded at a concentration of 3,333 cells per well in a 384-well non-adherent plate unless otherwise stated (Wako Chemicals, USA). They were cultured in EMEM supplemented with 10% FBS and 1% Penicillin-Streptomycin and allowed to grow for 48h before being introduced into the experimental device.

### Microfluidic device

The device design featured two main media channels adjacent to a central gel chamber (5 x 4.5 x 0.05 mm) with a partial wall covering 2/3 of the chamber’s height. A polymethylmethacrylate (PMMA) mold layout was created using Corel Draw (Corel Corporation USA) and was laser-cut using a CO2 laser (Universal Laser Systems Inc., USA). The partial wall was engraved on the surface of the PMMA mold. Bonding of PMMA layers was performed using 20% dichloromethane (DCM) and 80% Isopropyl Alcohol (IPA) at 60°C with a hot press. Polydimethylsiloxane (PDMS) (SYLGARD™ 184 Silicone Elastomer kit, Dow Corning, MI, USA) was prepared in a 10:1 ratio, poured into a mold, and cured overnight at 80°C. The devices were then cut from the mold and punctured with biopsy punchers to create a central hole and media reservoirs. The diameters of the punched holes were 4mm (media ports), 1mm (gel ports), and 2 mm (central hole). PDMS blocks were then autoclaved and treated for 90 seconds with air plasma (PE-25, Plasma Etch, NV, USA), bonded to coverslips, and kept at 70°C for 1 hour. Devices were then sterilized by UV prior to the experiment.

### Integrative vascular bed (iVas) formation

The iVas was created by self-assembly of endothelial cells (ECs) and FBs or human bone marrow-derived MSCs inside a fibrin gel. First, a fibrin gel solution containing ECs (8×10^6^ cells/ml) and stromal cells (1×10^6^ cells/ml) suspension and 6mg/ml fibrinogen gel solutions was injected into the gel channel of the device. The concentration of thrombin was adjusted so that when mixing 1:1 ratio with fibrinogen (6 mg/ml, Sigma-Aldrich), the gelation time reached approximately 2 min. The final thrombin concentration was approximately 2 U/ml, and the fibrin was 3 mg/ml. A total fibrin gel volume of 16 µL was injected into the central gel channels through one of the two inlets while tilting the device to ensure the even distribution of the gel. After removing the tip, the device was tapped gently to ensure the gel spread throughout the gel chamber while preventing it from entering the media channels. By injecting the gel slowly into its chamber, the gel spreads evenly over the chamber and spares the central hole, thus creating space for the tumor spheroid. The device was returned to a horizontal position for a gelation process, left at room temperature for 2 minutes, and placed at 37°C for 8 minutes. The central well can contain a thin layer of the gel due to initial injection, but this residual gel evaporates, creating an air-liquid interface that prevented the liquid from entering the hole. Next, the media was added to the media channels, 160 µL to one side and 40 µL to the other side, creating a pressure difference across the gel to stimulate the network formation. The media in the media channels was changed twice a day until spheroid insertion. A higher amount of media was applied to the opposite side of a previous channel with higher/lower media volume, which stimulated the network formation throughout the gel channel. Three days after the vascular bed seeding, the EC suspension (1×10^6^ cells/ml) was added to media channels. The devices were tilted for 30 minutes at 37°C after the EC suspension was added so that ECs could attach to the gel and form a monolayer. This procedure was then repeated on the opposite side. The monolayer serves to form openings in the media channel and connect to the vascular channel. Two days after the monolayer was formed, the device became fully perfusable and ready for spheroid insertion. The perfusion of the newly formed vasculature was confirmed using 10 kDa MW Cascade Blue Dextran solution (Thermo Fisher, USA).

### Spheroid insertion

Media channels were emptied on the day of spheroid insertion (Day 0), and 10 µL of 1× DPBS was pipetted into the central hole to break the surface tension. Media was then added to the media channels to create hydrostatic pressure within the central well, making the media fill up to the top of the central well. A tumor spheroid was transferred from the 384-well nonadherent plate to the central well using liquid-to-liquid contact via a wide-bore micropipette. The media at the media channel and central well were then removed by emptying the media in the media channels, and the central well was filled with a mixture of NaOH, rat tail collagen I (2 mg/mL, Enzo), fibrinogen (10 mg/ml), and thrombin (2 U/mL) in 1× DPBS. Subsequently, the EGM-2 media were mixed with EMEM with a ratio of 1:1 (EGM-2/EMEM 1:1) and used for media change.

### Drug-eluting shear-thinning hydrogel

STH was prepared according to a previous publication (9). Briefly, we prepared the stock solutions of 18% w/v type A porcine skin gelatin (Sigma Aldrich) and 9% w/v Laponite nanoclay (BYK Additives Ltd, TX, USA) using ultrapure water. The solid content was prepared by mixing gelatin/Laponite stock solution at 1:3 and 3:1 ratios to obtain shear-thinning biomaterials with 25% Laponite (STH25) and 75% Laponite (STH75), respectively. The total solid part of the gel solution is then combined with ultrapure water at a set weight to obtain a concentration of 6% (w/v) and mixed at 3000 rpm for 5 minutes using a SpeedMixer (DAC 150.1 FVZ, Germany). This mixing was repeated three times with a 5-minute pause between each round. The resulting Gelatin/Laponite STHs were stored at 4°C for 24 hours before use and then equilibrated at room temperature (25°C) for 60 minutes before experimentation.

### Loading of DOX into Shear-Thinning Hydrogels

To prepare DESTH, STH25 was mixed with an aqueous DOX solution (Oakwood Chemical, USA) to obtain a final DOX concentration of 100 µM in the hydrogel using a SpeedMixer at 3000 rpm for 5 minutes. Due to electrostatic charges, DOX binds to the Laponite nanoclay and is released into the TME under mildly acidic conditions (pH 5 and 7).

### Shear-thinning behavior and mechanical properties

The mechanical properties of the formulations were analyzed through injection force measurements. The Gelatin/Laponite STHs were placed in 3 cubic centimeter (cc) syringes and injected through 5.0 Fr microcatheters. Injection forces were measured using a mechanical tester (Instron 5943, Instron Int. Ltd., MA, USA) with the Bluehill version 3 software and a 100 N load cell, applying an injection rate of 100 mm/min.

### DOX Release Studies

To study DOX release, DESTHs were placed in buffers mimicking neutral and acidic physiological conditions, with pH 7.4 and pH 5.0. DOX release was monitored by measuring fluorescence at an excitation wavelength of 470 nm and an emission wavelength of 560 nm at various time intervals. Samples were taken over 10 days, and DOX fluorescence was quantified using a fluorometer to track release into the surrounding buffer solution.

### Chemoembolization treatment

One day after the spheroid insertion, devices were treated with DESTH and characterized for drug response. Depending on the group, STH25 with DOX (DESTH25) or drug-free STH25, the hydrogel was injected into one of the media channels via an angiographic catheter (Tip 5.0 Fr, Cook Beacon) using a syringe pump (World Precision Instruments, USA). The pump was set to apply a flow rate of 200 µL/minute. Once the hydrogel filled one media channel, we added 200 µL media (1:1 ratio of EGM-2 and EMEM media). After the chemoembolization, the pictures of the device were taken using a fluorescence microscope at different time points (1 h, 24 h, 48 h), capturing the spread of the DOX, which is autofluorescent in the red channel.

### Perfusion test

We perfused the vasculature networks with 10 kDa MW Cascade Blue Dextran or 2.0 µm fluorescently red polystyrene beads (Millipore Sigma) to check vascular network perfusability. For Dextran perfusion, the media channels were emptied, then 20 µL media containing 100 µg/ml Dextran was added to one side channel of the vascular network device. The other side was filled with 20 µL of the same media to immediately balance the hydrostatic pressure across the vascular bed. After the device was imaged, the media in the channels were changed at least two times to remove the Dextran from the device. For 2.0 µm bead perfusion, media channels had one side with 20 µL of media while a 5 cm-height media column in a 1ml syringe pressurized the other. Polystyrene beads (1:400 dilution from the stock solution in media) were injected into the media column and dragged into the vasculatures by the flow generated by the hydrostatic pressure gradient.

### Flow experiment setup

For the continuous flow experiment for evaluating embolic agents, two different chemotherapy-free embolization materials were used – STH25 and commercially available TACE Embozene^TM^ Microspheres 40µm (Varian, CA, USA). The embolization of one of the media channels was carried out 1 day after the insertion of a HepG2 spheroid into the device. The 1ml syringe columns tightly fit into three media inlets. The fourth media reservoir, on the non-embolized media channel, was connected to the syringe by a custom-made connector that had a tight fit to the PDMS chip on one side and a luer lock on the other side, which connected to a tubing routed to a syringe. The syringe was then installed to the pump and set on the withdrawal mode with a 1 µL/min flow rate. The media were prepared by mixing 1:1 EGM-2 and EMEM media. We topped up the media reservoir daily. The experimental results were collected 4 days after the embolization. Moreover, STH75 was employed to seal any potential leakage from the central hole before introducing the media.

### Testing the blocking of the flows using acellular devices

To evaluate interstitial flows across acellular fibrin gels to compare the blocking efficacy of different embolic agents, the gel channel of a microfluidic chip was first filled with fibrin gels. Its media channels were then pressurized by a 5 mm H_2_O hydrostatic pressure, with or without the presence of an embolic agent in the media channel (STH25 or Embozene^TM^).

### Image acquisition, image processing, and analysis

Live images were taken using an Echo Revolution microscope (Echo, CA, USA) with a built-in 1.25x lens. Immunofluorescence images were acquired on a Zeiss LSM710 confocal microscope (Zeiss, Germany). The filter sets that were used for the microscope were DAPI (EX/EM = 380/450 nm), FITC (EX/EM = 470/525 nm), TxRED (EX/EM = 560/630 nm), CY5 (EX/EM = 630/700 nm).

To measure the microvascular’s mean diameters, the Dextran channel of an image of a perfusable area of the microvasculature on day 0 was converted into a mask (**Fig. S1**), and MATLAB Reaver was used to calculate the mean diameters (30). For the vascular morphology analysis in the flow experiment, we combined built-in ImageJ machine learning-based segmentation and MATLAB to extract vascular density, number of branch points (points in the network where a single vessel bifurcates into two or more vessels), number of endpoints, and the maximum branch length. First, the mask of vasculatures was obtained using Weka trainable segmentation plugins. Vascular density was measured by dividing the vasculature’s surface area by the gel channel’s total area. The number of endpoints, presenting the discontinuity of the networks, was obtained using the Analyze Skeleton plugin function; the number of branch points, presenting the network connectivity, and total length of all branches were valuated using MATLAB’s REAVER plugins (30, 31).

### ELISA

Albumin and VEGF-A ELISA kits (Thermo Fisher, USA) were used according to the manufacturer’s protocol. The fluorescence intensity readouts of the plates were performed using a microplate reader (Varioskan LUX, Thermo Fisher), with excitation at 560 nm and emission at 590 nm.

### Measurement of cytokines in media

The media from the device media channels were collected for the Human Cytokine/Chemokine 48-Plex Discovery Assay® Luminex assay performed by Eve Technologies (Alberta, Canada). Cytokine analysis was performed using media from devices treated with DESTH, STH25 gel only, and control devices without spheroid or with HepG2 tumor spheroid condition media at 48h. The lowest detectable concentration of the standard curve replaced all out-of-range data of each cytokine in the panel. Data were plotted in a log10 scale of concentration values normalized to the control device with a HepG2 spheroid. The Luminex assay data were obtained from at least 3 devices using pooled culture media.

### Immunofluorescence staining

The cell culture media was discarded first, and then, we rinsed the devices with 1× DPBS. Next, the samples were fixed with 4% paraformaldehyde (PFA, Thermo Fisher Scientific, MA, USA) for 20 minutes. After the fixation, the devices were washed thrice with 1× DPBS. Then, DPBS with 0.1% Triton X-100 (Thermo Fisher Scientific, MA, USA) was applied to the device on a shaker at room temperature (RT) for 15 minutes. Following this, cell blocking was carried out by administering a solution of 5% w/v BSA (Millipore Sigma, MO, USA) dissolved in 1× DPBS for one hour at RT. Mouse anti-human EpCAM (IgG2b, kappa, 9C4) (Biolegend) was used for the immunostaining process. The antibodies, diluted to a ratio of 1:50 in DPBS with 0.1% BSA, were applied to the devices and left to incubate overnight at a temperature of 4°C. The following day, the devices underwent two washes using a wash buffer composed of DPBS with 0.1% BSA.

Secondary immunostaining was performed by applying antibodies at a dilution of 1:200, which were conjugated with Alexa Fluor®594 goat anti-mouse IgG (H+L) (Invitrogen, Carlsbad, CA, USA), and allowing them to incubate on the devices overnight at 4°C. A subsequent triple wash using DPBS with 0.1% Triton X-100 was conducted. The process concluded with DAPI counterstaining and an additional wash using DPBS 1×.

### Viability assays

Live-dead staining was performed using Calcein AM and Ethd-1 double staining (Fisher Scientific, MA, USA). HepG2 spheroids were treated with 0, 0.1, 1, 10, and 100 µM DOX for 24 hours. Then, we washed the spheroids using DPBS and added Calcein AM and Ethd-1 diluted at working concentrations (2 µM and 4 µM) in DPBS for 30 minutes.

PrestoBlue viability assays were performed according to the manufacturer’s protocol. Tumor spheroids were seeded at a concentration of 3333 cells/well for 3 days and treated for 24h. The readout was performed using a Varioskan LUX microplate reader (excitation/emission wavelengths: 560 nm/590 nm).

### Simulation of DOX distribution

A computational model based on COMSOL Multiphysics was created to investigate the diffusion of the DOX molecule within the fibrin bed in the device, which includes a circular tumor region with a 2 mm diameter. Given that the dimensions of the gel region outside of the hole in a device are 0.5 mm in height, 5 mm in width, and 4.5 mm in length, with the height being much less than the width and length, the simulation was assumed to be two-dimensional. We used the “Transport of Diluted Species in Porous Media” module for saturated porous media (no gas phase). The mass balance equations for this model, with the assumption of Newtonian, isothermal, and incompressible fluids, are as follows (32):

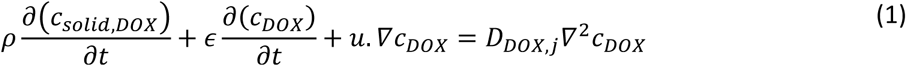

Here, *ρ* (kg/m^3^) refers to the bulk density of fibrin, ɛ is the porosity or the liquid volume fraction in the gel channel, *c*_*DOX*_ (µM) is the concentration of DOX in the media, *c*_*solid*,*DOX*_ the concentration in µM of DOX absorbed in the solid part of the gel bed, **u** is the velocity field (SI unit: m/s) and *D*_*DOX*_ the diffusion coefficient of DOX (m^2^/s). We incorporated the effects of porosity and tortuosity of the medium in different regions; j = 1 indicates the compartment inside the gel channel and outside the central hole, or j=2 indicates the region inside the central hole. The parameters of the simulations are as follows: *ε* =0.25, *D*_*DOX*,*j*_ ∼= 10^-9^ m^2^/s (for both j=1 and 2), estimated based on:

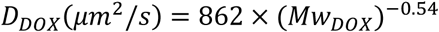

With DOX molecular weight *Mw*_*DOX*_=0.54 kDa and the fibrin bed assuming an extracellular matrix similar to published literature (35). The term 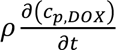 corresponds to changes in the concentration within the fibrin matrix due to absorption or desorption. Since we assumed negligible absorption or desorption of DOX in the matrix, and hence 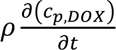 was neglected. Because the gel blocks the flow going across the vascular bed, *u*. *▽c*_*DOX*_=0. The equation (1) can be rewritten as follows:

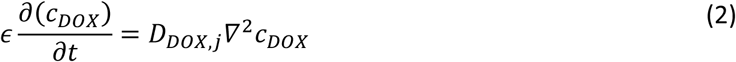

For boundary conditions, *c*_*DOX*_ (t=0) =0 µM, with j = 1 or 2. The boundaries of the simulated domain were considered to be impermeable and non-slipping. At the right of the fibrin bed (x = 5 mm), *c*_*DOX*_ =100 µM, while the left side (x = 0 mm) has *c*_*DOX*_ =0 µM, respectively. The computational model was divided into ∼28000 triangular meshes to ensure grid independence.

To compare the simulation results with the experimentally acquired DOX fluorescence signals, we plotted the total concentration of DOX at a location outside the central well = *c*_*total*,*outside*_ = *c*_*DOX*_. ℎ, with ℎ is the height of the chamber. Concentrations of DOX at different time points (t=0, 1, 24, 48 hours) were plotted in all positions in the gel channel. Considering the height of the gel in the central hole is higher than the gel in the gel channel outside the central well, we need to multiply ℎ^∗^ to the *c*_*total*_ in the central well by a factor *w*: *c*_*total*, *inside*_ = *c*_*DOX*_. ℎ^∗^with ℎ^∗^ is the maximum height where the fluorescence signal can be detected in the column. We can estimate ℎ^∗^ by computing it from experimental measurements:

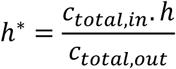

To quantify the ratio between *c*_*total*,*in*_ and *c*_*total*,*out*_, we measured fluorescence signal intensity in the TxRED channel (Ex/EM 560/630 nm) of the device at 48h at the equilibrium in a 200 µm × 200 µm region of interest located in either the central well region or the gel channel outside the central well, yielding in 5416 fluorescence units with standard deviation (SD)= 468 and 4251 fluorescence units with SD= 539 respectively.

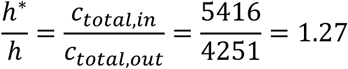

Therefore, ℎ^∗^ = 637 µm. A Python program was used to convert the cartesian coordinates of DOX diffusion simulations into polar coordinates. Positions within the central well were those with a distance < or = to 1mm from the position x=2.5, y=2.25 mm.

### Statistical analysis

Unpaired Student’s t-tests were used to compare two independent groups following normality tests. A one-tailed t-test was used to compare albumin levels in devices treated with DESTH and control devices, as a decrease in albumin was decreased following TACE treatment (33). As observed in flow experiments, paired Student’s t-tests were employed when the data sets were paired. One-way ANOVA tests were utilized for analyses involving multiple groups, with post hoc evaluations conducted using Dunnett’s test for comparisons to a control group or Tukey’s tests for intergroup comparisons. Non-significant comparisons are not reported.

Analytical procedures were executed utilizing Prism 7 software (Dotmatics, USA) or Python. Each analysis was based on the mean values derived from n≥3 microfluidic devices, with each device corresponding to a distinct experiment. Significance thresholds were established at * p<0.05, ** p<0.01, *** p<0.001, and **** p<0.001.

## RESULTS

### Establishment of a vascularized liver tumor model

To develop our vascularized liver tumor model, we first created iVas by seeded HUVECs with stromal cells such as FBs or MSCs within fibrin gels into the microfluidic chamber that has an open-top well (**Fig. 1A**) and a tumor spheroid was formed using low-adhesion 384-well plates separately (**Fig. 1B**). Using the meniscus trap technique (22), an empty well was created within the fibrin gel solution after deposition by surface tension (**Fig. 1C**). Media in the media channel did not fill the central hole channel after fibrin was solidified. Therefore, self-assembled microvasculatures were later formed around the well without infiltrating it. Within this setup, endothelial cells, aided by paracrine signaling from either FBs or MSCs, organized into functional vascular networks, forming the iVas. After the vascular networks were formed, the tumor spheroid was transferred from a well plate into the central hole, flanked by fibrin and collagen extracellular matrix, and surrounded by vasculatures. One day after co-culture, the networks were perfusable in the area surrounding the tumor spheroid, shown by blue Dextran perfusion (**Fig. 1D-E**). Vascular beds created by seeding ECs and MSCs formed more narrow, not perfusable vessels (**Fig. S2**). Therefore, in this study, we focused on vascular bed models created by ECs/FBs co-culture. The histogram of the diameters of the microvascular bed showed that the *in vitro* microvasculature (Mean diameter: 45 µm, SD: 9.9 µm) has a similar size to HCC capillary *in vivo*, which has the smallest diameters between 30-40 µm see **Fig. 1F** (34). VEGF ELISA analysis showed the secretion of VEGF by tumor spheroid, explaining molecular interactions between the tumor spheroids and the vascular networks supporting vascular stability (**Fig. S3**).

**Figure 1:**
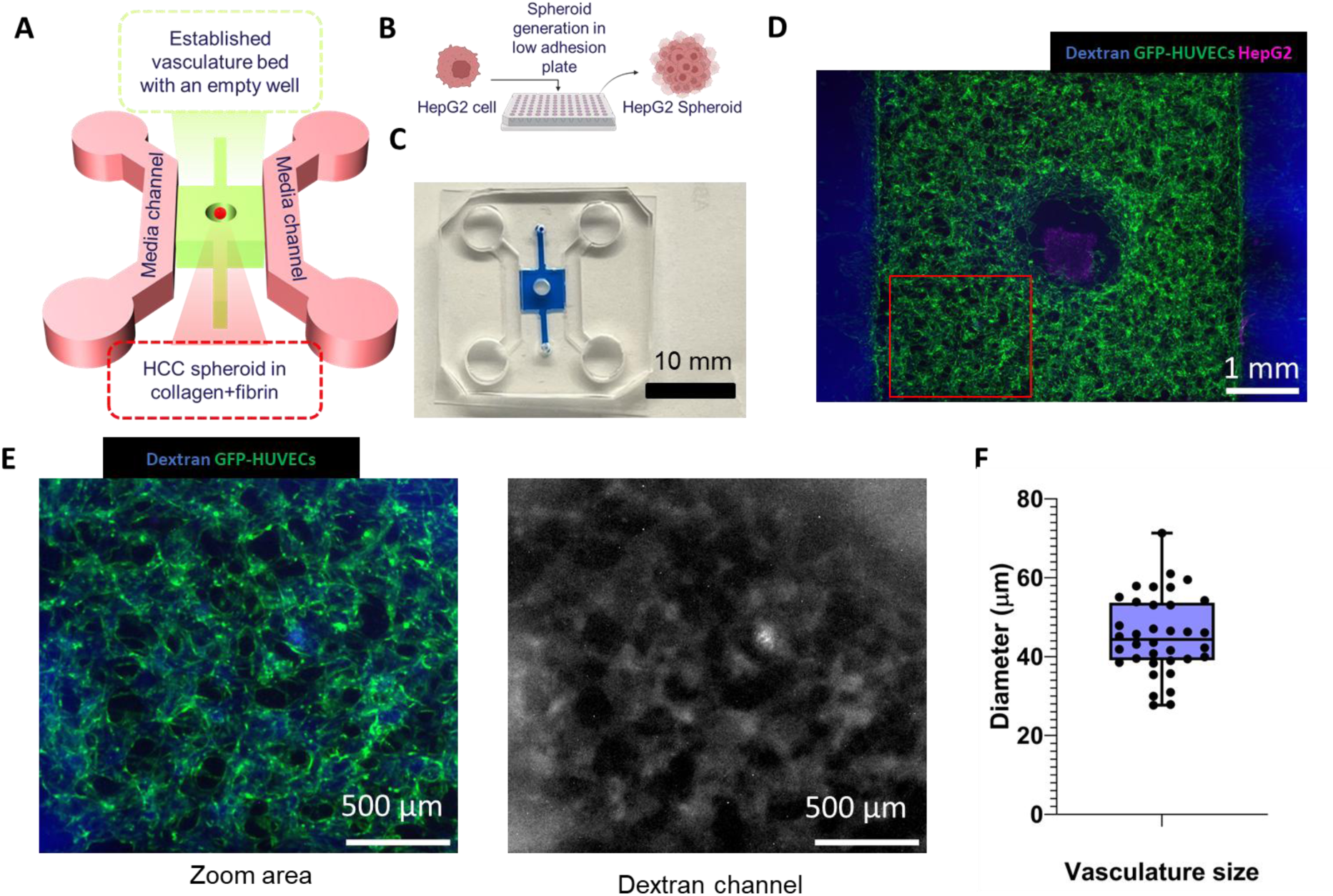
Establishment of a vascularized liver tumor model. A) Schematic presentation of a microfluidic device with a central empty well to insert a HepG2 spheroid. B) Schematic presentation of tumor spheroid creation using a low-adhesion well plate. C) Representative image of a microfluidic chip with the gel channel filled with a blue dye solution to visualize the gel channel. At the center of the vascular bed, an empty well is created by trapping the meniscus within the microfluidic chamber. D) Vasculatures are formed by seeding human umbilical vein endothelial cells (ECs) and human lung fibroblasts (FBs). The functional vasculature is perfused with 10KDa Dextran. E) Zoom-in view demonstrating Dextran within microvasculatures: the left shows all channels, and the right displays only the blue channel (Dextran). F) Mean microvasculature diameters measured in a batch of 36 devices.

### Optimization of DOX concentration and rheological properties of DESTH

To make DESTH specific to this setup, we first needed to optimize the concentration of DOX used for the drug release. When spheroids containing ∼1.3×10^4^ HepG2 cells per spheroid were initially seeded, they exhibited limited cohesion (**Fig. S4**). Consequently, we chose spheroids containing ∼3×10^3^ cells each, which displayed a more compact structure. Next, to optimize the drug concentration for treating HepG2 spheroids, we treated these spheroids with free DOX across a concentration range of 0, 0.1, 1, 10, 100 µM for 24 hours. Live-dead staining revealed that tumor cell death was most pronounced at 100 µM DOX concentration (**Fig. 2Ai-v**). Metabolic assays also confirmed the effectiveness of 100 µM DOX compared to lower concentrations (**Fig. 2B**). Consequently, we chose 100 µM DOX for the formulation of DESTH.

**Figure 2:**
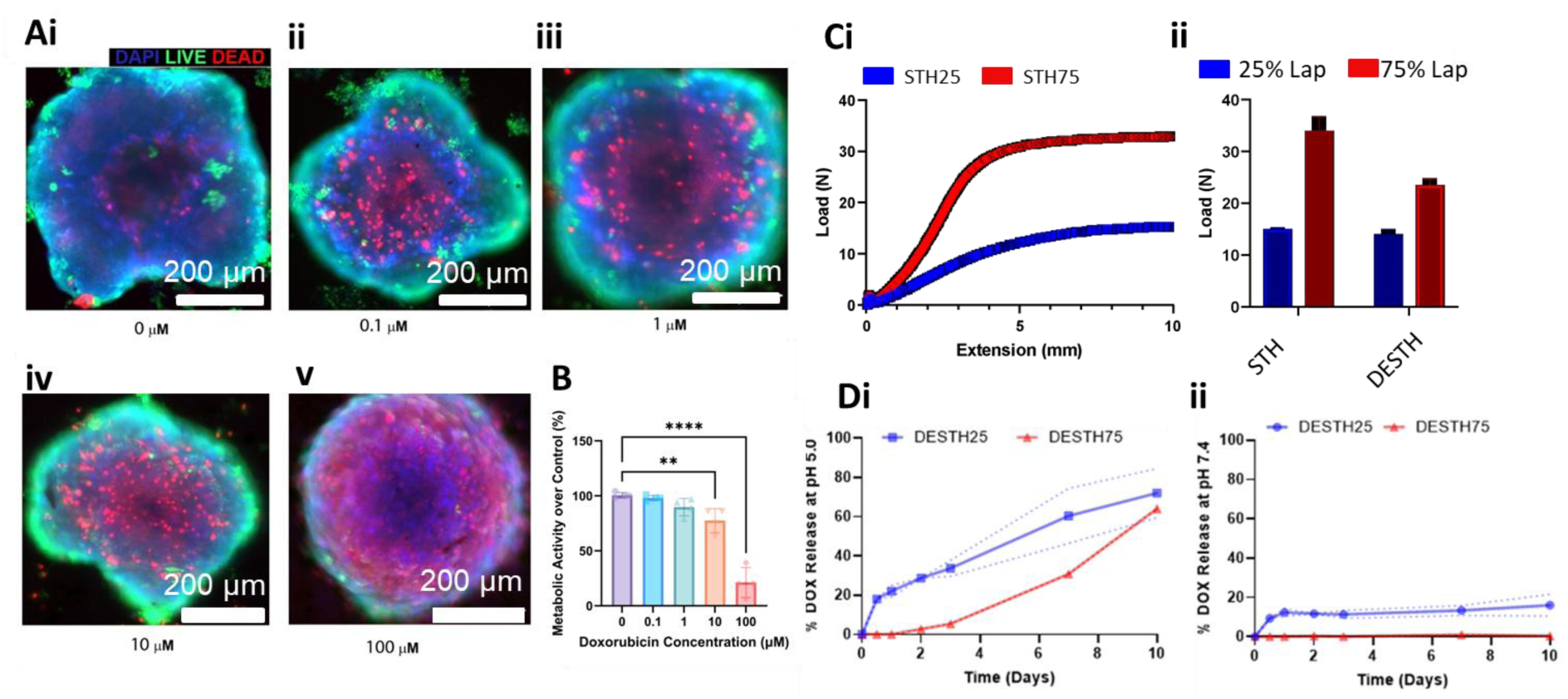
Drug-eluting shear thinning hydrogel (DESTH) optimization. Ai-v) HepG2 response to different concentrations of free Doxorubicin (DOX) (0, 0.1, 1, 10, and 100µM, respectively). B) Metabolic activity of HepG2 spheroid associated with each free DOX concentration. C) Mechanical properties of DESTH. C) Injection force measurements of STH. Ci) Extension and injection load relationship of STH25 and STH75. Cii) Maximal injection load comparison of STH25 and STH75 with or without DOX. D) DOX release profile of DESTH25 (STH25+DOX) and DESTH75 (STH75+DOX), at Di) pH5 and Dii) pH7.4. Statistical tests for 2B are one-way ANOVA with Dunnett’s tests, ** p<0.01, and **** p<0.0001.

We developed STHs by mixing Laponite nanoclay with gelatin, following our previously established protocol (9). Injectability testing indicated that STH25, without DOX encapsulation, required around 15N of force with a 3 cc/5 Fr syringe setup, while STH75, with the higher Laponite concentration, needed 34N and STH75 with DOX encapsulation needed 23N, approximately, see Fig. 2Ci and ii. This aligns with our previous studies suggesting that nanocomposites with lower Laponite-to-gelatin ratios retain gelatin’s viscoelastic properties, while hydrogels with a higher Laponite-to-gelatin ratio exhibit increased viscosity and more pronounced shear-thinning behavior (9). Finally, hydrogels with a lower Laponite concentration (STH25) released roughly 35% of the loaded DOX within three days at an acidic pH, while those with higher Laponite content, such as STH75, released less than 5%, see Fig. 2Di. After three days in an acidic environment, DOX release significantly increased, reaching as high as 70% over 10 days. At a neutral pH, all STH formulations released less than 10% of DOX within the first three days, with no notable increase over extended incubation time (**Fig. 2Dii**). Therefore, we chose STH25 as the material for DESTH as its drug release was faster during the experimental window.

The injection of DESTH into vascularized liver tumor spheroid models resulted in the death of tumor cells and the regression of the vasculature. We performed embolization of a microfluidic chip using a catheter loaded with DESTH (**Fig. 3A**). Prior to embolization, the vasculatures surrounding the liver tumor were perfusable, shown by blue Dextran (**Fig. 3Bi, ii**). Following embolization, one side channel was filled with DESTH (**Fig. 3Biii**), thereby blocking the outlet of the microvasculature networks. Consequently, microvasculature networks were no longer perfusable (**Fig. 3Biv**). At 24h after applying DESTH, the vasculatures began to regress at the region close to the DESTH-filled channel (**Fig. 3Bv**). At 48h after the DESTH application, the device exhibited narrow and disrupted vasculatures (**Fig. 3Bvi**). Live-dead staining of tumor spheroid at 48 hours post-treatment revealed that the majority of HepG2 tumor spheroids were dead in the DESTH condition (**Fig. 3C**). Given the red autofluorescence red of DOX, the diffusion of DOX from the DESTH to the liver tumor models was characterized at 24 and 48 hours. The resulting images showed the diffusion of DOX into the gel channel and the central hole (**Fig. 3D**). Our COMSOL model also shows that DOX from the DESTH reaches the central channel after 1 hour and is at equilibrium after 24 and 48 hours (**Fig. 3E**), helping the study of the distribution of drugs within the TME.

**Figure 3:**
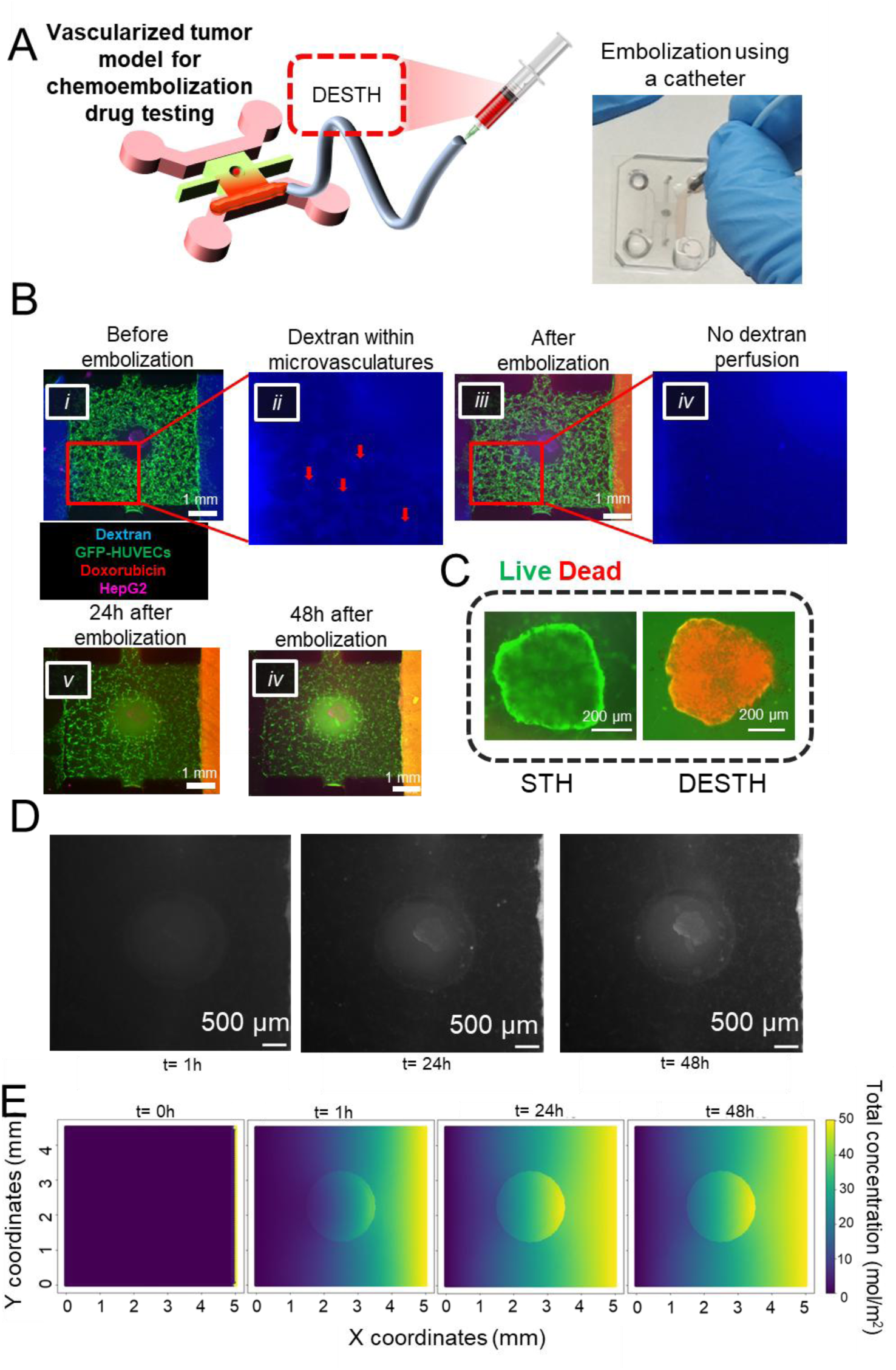
In vitro chemoembolization of a vascularized liver tumor model. A) Schematic illustration of DESTH treatment in a vascularized liver tumor model. Drug-eluting shear-thinning hydrogel (DESTH) containing DOX is injected into a vascularized liver tumor model via a catheter. B) Fluorescence images of vascularized liver tumor models before and after treatment with DESTH. i) The vascularized tumor model prior to treatment shows perfusable microvasculatures filled with Dextran. Red arrows indicate microvasculatures filled with Dextrans. ii) A zoom image from Bi showing the Dextran channel. iii) Microvascular bed right after treatment with DESTH. iv) A zoom image from Biii showing the microvasculatures are not perfusable with Dextran. v) 24h and vi) 48h post-treatment with DESTH, showing regression of vasculatures. D) Distribution of DOX within the gel chamber over 1, 24 and 48h. E) Simulation of DOX distribution in the device at t=0, 1, 24, and 48 hours.

To evaluate responses to chemoembolization, live/dead staining was performed on devices treated with control, drug-free STH25, and DESTH25 for 96 hours (**Fig. 4A**). In comparison with the control, a subset of tumor cells exhibited a state of necrosis in devices treated with drug-free STH. In contrast, the live dead staining revealed low viability of HepG2 cells when the device was treated with DESTH. EpCAM immunostaining of devices, both untreated and treated with STH25 or DESTH25 (**Fig. 4B**), demonstrated that HepG2 cells retained EpCAM expression in drug-free embolization treatment. However, EpCAM expression immunofluorescence signals were undetectable after the DESTH25 treatment. Therefore, the observed loss of EpCAM expression can be attributed to the release of DOX from DESTH25, which is consistent with a previous finding that EpCAM+ cell lines, such as HepG2, exhibit a decrease in EpCAM expression after treatment with DOX (35). Moreover, analysis of pooled culture media after 48 hours of treatment reveals the upregulated inflammatory cytokines in DESTH-treated devices compared to untreated samples, such as 59-fold increase of IL-1α, 15-fold increase of GM-CSF, 7.7-fold increase of VEGF, 5.8-fold increase of IL-4, and 4-fold increase of IL-1β (**Fig. 4C**). Importantly, the expression levels of IL-4 and VEGF were also observed in HCC patients treated with TACE (36, 37). The increase of proinflammatory cytokines such as GM-CSF, IL-1β can be explained by DOX-induced accumulation of reactive oxygen species in HepG2 cells (38). These cytokines are relevant activators of the immune system and angiogenesis. Moreover, albumin secretion in devices treated with DESTH was found to be lower in comparison to the control devices, indicating a reduced level of HCC activity in the treated devices (**Fig. 4C and 4D**). This finding is consistent with a prior study showing decreased HCC patient’s serum albumin levels after TACE treatment (33).

**Figure 4:**
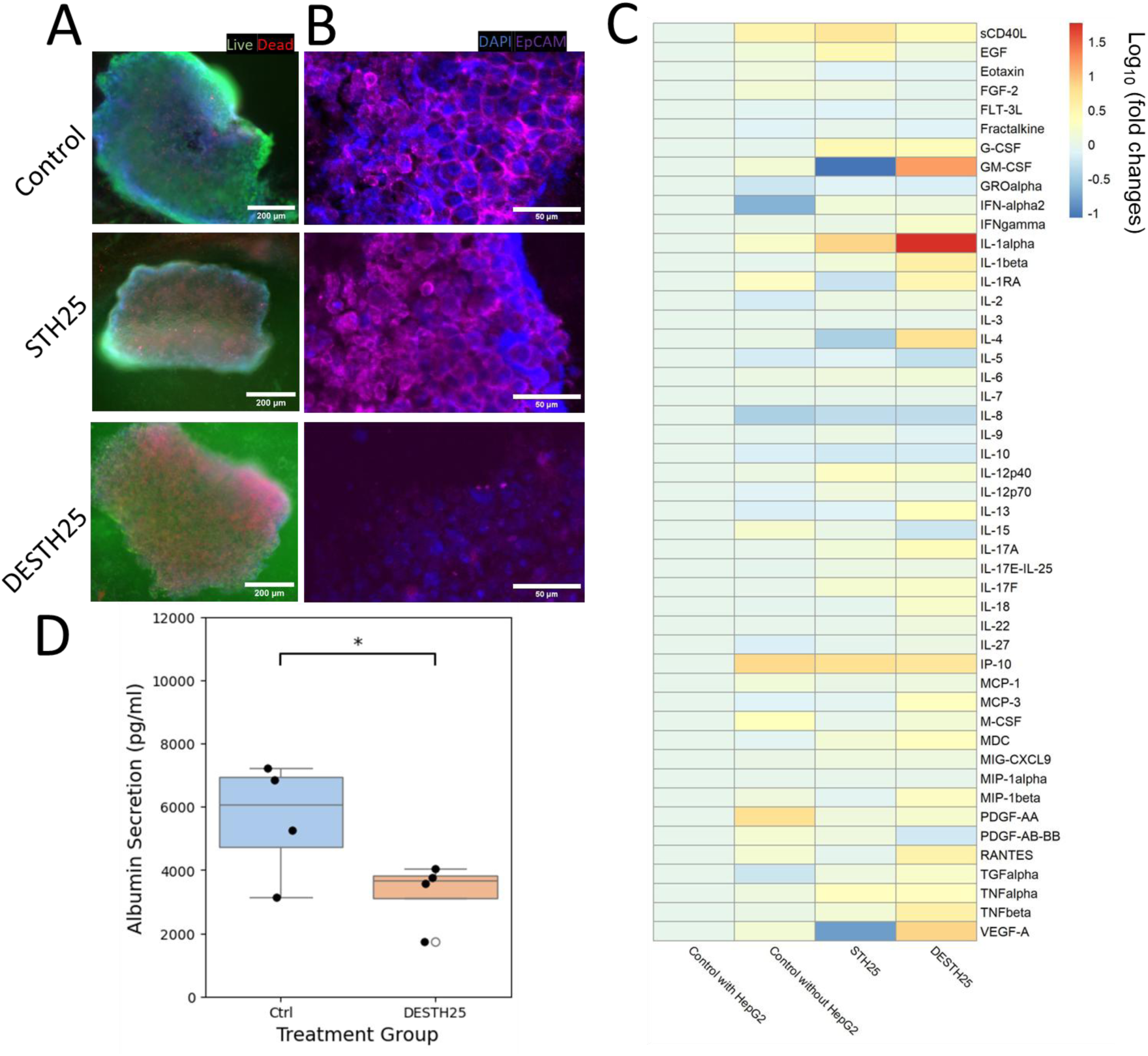
Characterization of cellular and cytokine responses to DESTH. A) Live/dead staining of devices 96 hours after embolization with: i) control and ii) STH25, iii) DESTH25 conditions. B) EpCAM immunostaining of HepG2 tumor spheroid inside vascular bed devices under various treatments after 96h. C) Cytokine analysis of control devices without a tumor spheroid, control devices with a HepG2 spheroid, devices treated with DESTH, STH25 gel only, control device without spheroid or HepG2 tumor spheroid condition media at 48h. Data are plotted in a log e scale of concentration values normalized to the control device with a HepG2 spheroid. D) Comparison of albumin secretion between the control and DESTH25-treated devices. Statistical tests are one-tailed Student’s t-test.

### Microfluidic flow allows capture of embolization effects of various embolic agents

The verification of flow blockage was conducted by applying a colored dye solution and hydrostatic pressure to an acellular fibrin gel on a chip. The results demonstrated that the device treated with STH25 achieved complete flow blockage, while the Embozene^TM^-treated device exhibited only partial blockage (**Fig. 5A**). To emulate the flow from the portal vein and hepatic artery to the hepatic vein (**Fig. 5B**), we performed a flow experiment using the vascularized liver model. Each setup consisted of 2 coupled devices: one treated with an embolic agent, either STH25 gel or Embozene^TM^, and the other is the untreated control device (**Fig.5C**). During the flow experiment, vasculatures were compared between the two devices: one embolized and one control, which has 5 mm H_2_O hydrostatic pressure (∼0.007 psi) generated by columns of media on all three inlets, while the 4^th^ outlet has a negative pressure caused by an aspirating syringe.

**Figure 5:**
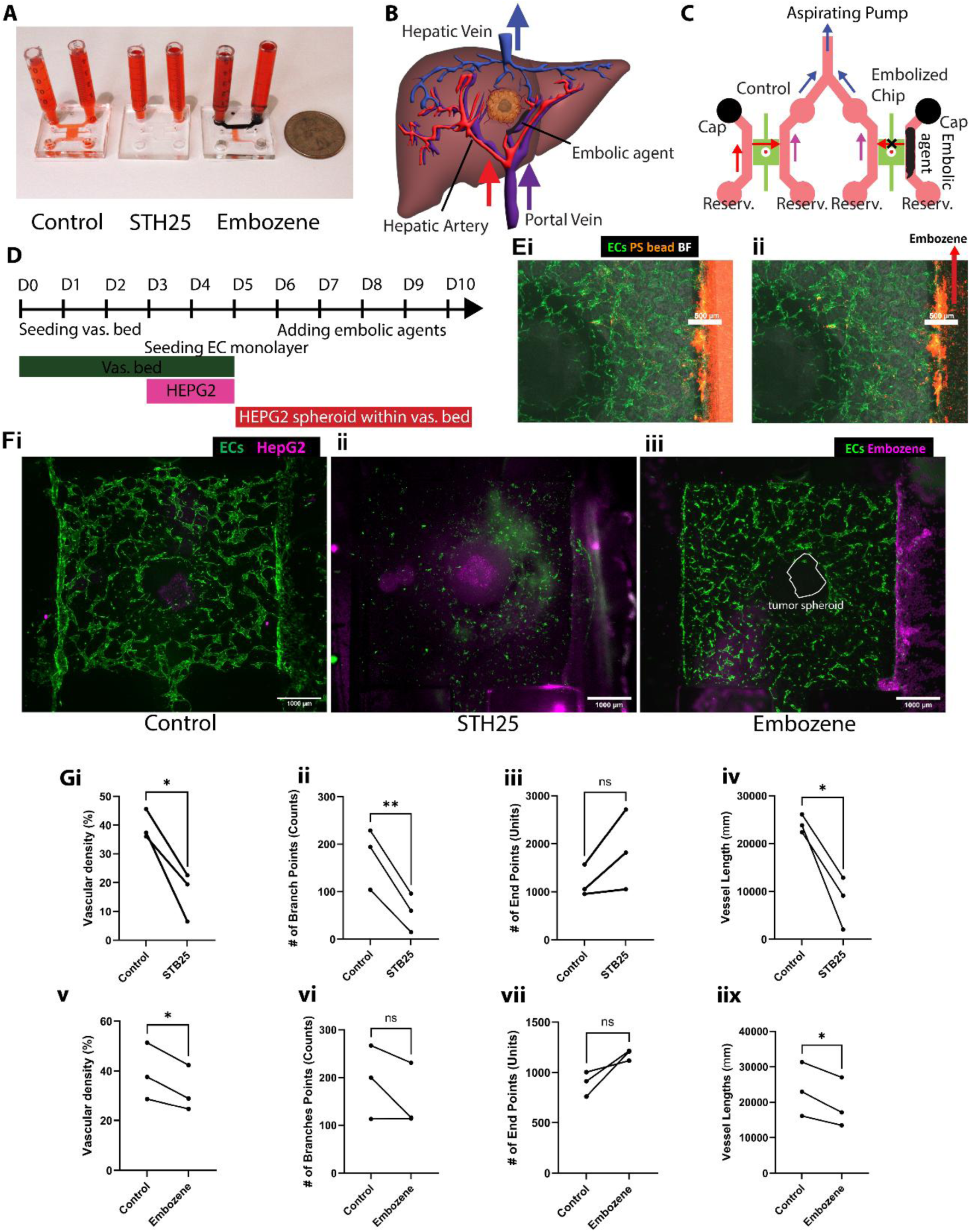
Evaluation of embolization effects in a vascularized liver tumor model under perfusion. A) Transport of red dyes by interstitial flow across cell-free fibrin gel within a microfluidic chamber when the side channel is not blocked or blocked with STH25 or Embozene^TM^beads. B) Visual representation of liver vasculature system. The liver vasculatures have two inlets: the hepatic artery and portal vein, and one outlet, the hepatic vein. Liver tumor embolization happens on the artery side, blocking blood from entering the tumor through the liver artery branches. C) Schematic presentation of fluid flows within the fluidic system in the flow experiment. Each vascular bed has two inlets and one outlet. Two column reservoirs are attached to the chip, one on each side of the vascular bed. The media is drawn through an outlet into a syringe by a pump. When the chip is embolized on one side, there is no flow across the vascular bed, cutting the supply to the microvascular networks and the tumor spheroid. For the control chip, the media continues to flow through the microvascular bed, sustaining the vasculatures and the tumor spheroid over 4 days of flow experiment. D) Timeline of the experiment. D) Perfusability of the vascular bed before and after Embozene^TM^ treatment. Ei) Fluorescent polystyrene (PS) bead (orange) perfusion into vascular bed (green). Eii) Embozene beads, shown in bright-field (BF) images, are perfused to the right side channel. F) Images of the vascular bed device on day 4 after starting the flow experiment in different conditions: i) Control device, ii) STH25 gel, iii) Embozene^TM^. G) Vascular morphology analysis. i-iv) Vascular density, number of branch points and endpoints, and maximum vessel lengths of the STH25-treated devices and their controls. v-iix) Vascular morphology parameters of devices treated with Embozene^TM^ compared to their controls. P-values were obtained using paired Student’s t-tests.

A HepG2 tumor spheroid was inserted into the vascular bed 24h before the treatment with embolic agents and flow experiment (**Fig. 5D**). To demonstrate embolization blockage, small-size fluorescence beads (2 µm) were perfused into the vasculatures by applying hydrostatic pressure at the side channel (Video S1). The same side channel was subsequently filled with 40-micrometer Embozene^TM^ beads, resulting in partial blockage of the flow (**Fig. 5E i and ii**). Indeed, we found that the untreated control devices, which received flows across the perfusable vasculature throughout 4 days of the experiment, still had an interconnected structure with high density (**Fig. 5Fi**). In contrast, the devices treated with STH25 exhibited a significant regression (**Fig. 5Fii**) and the Embozene^TM^-treated devices demonstrated a regression to a lesser extent (**Fig. 5Fiii**). Quantitative vascular morphology assessment demonstrated significant reduction of network density, connected branch points, and overall length in devices treated with STH25 compared to the control (**Fig. 5Gi-iv**). In contrast, devices treated with Embozene^TM^ showed less change in vasculature structure than STH25 compared to the control, particularly in connectivity, as indicated by the number of branch points (**Fig. 5Gv-viii**). These findings collectively demonstrated that the complete obstruction of flow through the microvasculature by STH25 resulted in the collapse of the vasculature.

## DISCUSSION

In our study, we engineered a vascularized liver tumor model to develop and validate novel embolic agents. We built a fully perfusable vascular bed surrounding a central well accommodating liver tumor engraftment. This design allows the supply of nutrients from the side channels to the central well via the microvascular networks for the survival and growth of the tumor cells.

The dimensions of our *in vitro* vascular networks closely resemble *in vivo* HCC capillaries, recapitulating local HCC microcirculation. Unlike a healthy liver with sinusoid networks, HCC remodels the vascular networks and receives blood through hepatic arteries and capillaries (39). In our model, vasculature diameters ranged between 27 to 72 µm, mean diameter = 45 ± 9.9 µm. These *in vitro* microvessels are similar to the smallest visible vessels in human HCC microvasculature network revealed by X-ray phase-computed tomography, which ranged between 30-40 µm (34).

Developing preclinical models that accurately mimic human pathophysiology is key for finding new therapies, given the high failure rates in drug development, often attributed to the limitations of conventional cell culture and animal models (40, 41). Furthermore, the implementation of 3R principles of Replacement, Reduction and Refinement of animal use in research, adopted in the European Union Directive 2010/63/EU, is prompting the scientific community toward utilization of alternative methods (42). Vascularized cancer models address this issue by reconstructing a multicellular architecture and physiologically relevant tumor microenvironment *in vitro*, and enabling vascular perfusion that better reflects *in vivo* conditions (18, 43). As TACE requires precise, catheter-based delivery of chemotherapy into the hepatic artery near a liver tumor, creating a microphysiological system recapitulating this process allows for the development of new TACE compounds. STH has the unique ability to lower viscosity under shear stress and then recover once the stress is removed, facilitating smooth injection through needles and catheters. To assess the efficacy of the newly developed DESTH, we proposed a microfluidic-based framework designed to emulate TACE process in a physiologically relevant model. The study was divided into two main parts: 1) static chemoembolization, which tests the effect of DESTH on tumor cells and vasculatures 48 hours after treatment, and 2) a flow experiment, which evaluates the effect of embolic agents on vasculatures over 96 hours post-treatment. First, we used this platform to evaluate the response of the HCC vasculatures to DOX released from DESTH, tracking drug distribution over time and the time-dependent response of the human tissue model. STH25, with a lower Laponite content, exhibited better injectability and faster drug release at low pH than STH75, with a higher Laponite content. Treatment with DESTH made by mixing STH25 and DOX on a vascularized tumor model results in tumor cell death and vasculature regression within the tumor microenvironment. Our findings suggest that increased VEGF and IL-4 and decreased albumin levels are markers for assessing response to chemoembolization in our in vitro model. These observations agree with the published data from HCC patients treated with TACE (33, 36, 37). Second, to distinguish the effects of embolization and chemotherapy on the model, we performed flow experiments using our vascularized liver tumor spheroid models to assess the efficacy of different embolic agents. In the absence of the flows inside the microvascular bed, the vasculature bed regresses and becomes smaller and shorter compared to the control vasculatures. Moreover, different embolic agents result in different vascular morphologies, highlighting the potential use of these models to optimize the embolic agents.

The use of vascularized liver tumor models in our flow experiments provides a closer representation of the *in vivo* tumor microenvironment than conventional cell cultures or animal models (44). In particular, two recent publications from Bonanini et al. proposed novel vascularized liver models: one is based on grafting primary human hepatocyte spheroid into a vascular bed created by endothelial angiogenesis sprouting and the other is based on vasculogenesis of endothelial cells when co-cultured with liver cells such as hepatocytes and stellate cells (45, 46). Compared to these studies, our system is based on engraftment of tumor spheroid on vasculogenesis-based microvascular bed with a central well for spheroid transfers. It ensures consistent and robust perfusability of the vasculatures surrounding the engrafted tumor spheroid and enables the modeling of liver embolization therapy. As the models incorporate the vascularized tumor spheroids and controlled flow conditions, they allow for the study of vascular regression caused by embolic agents. This model allows the distinction of embolization efficacy of various embolic agents through real-time characterization of drug distribution and vascular morphology. Therefore, the future development of novel therapeutic strategies that combine targeting the tumor and its vasculatures, as well as the study of drug delivery in the TME, could be done using physiological models.

In the future, we will need to improve our model by adding other liver tumor stromal cells such as cancer-associated fibroblasts, hepatic stellate cells, healthy hepatocytes, and other immune cells (Kupffer cells, T-cells) to create a model mimicking closely to the liver TME (47). Another consideration is to improve the model for assessing embolization-related ischemia effects. This could be done using a hypoxic incubator and oxygenated media carrying red blood cells to mimic oxygen diffusion from the media to the tissue model. Finally, long-term effects and potential adverse reactions need to be evaluated over a longer period with optimized media to ensure the safety and durability of the embolic agents.

## CONCLUSION

In conclusion, our study created a novel *in vitro* model that mimics the liver cancer capillary system for embolic agent testing. It demonstrates the efficacy of injecting DESTH in inducing tumor cell death, triggering proinflammatory cytokine release, and decreasing albumin secretion in vascularized liver tumor spheroid models. Evaluating different embolic agents under flow conditions offers insights into vessel regression, paving the way for testing different embolization agents and strategies. By bridging the gap between animal models and human physiology, this model offers novel indications of cancer treatment efficacy by investigating cellular and cytokine responses to therapy. We anticipate that the findings from this study will contribute to the refinement and development of embolization techniques, ultimately benefiting patients with various vascular diseases, particularly liver tumors.

## CONFLICT OF INTEREST

No conflict of interest

## Acknowledgement

The authors gratefully acknowledge funding from the National Institutes of Health (R21EB034423, R01AI183529, R01AR074234, UH3TR003148, R34NS126032, R01GM126571, CA257558, and DK130566). N.F. appreciates the postdoctoral fellowship from the National Science and Engineering Research Council (NSERC). We thank Dr. Marvin Mecwan for his help with the experimentation.

## SUPPLEMENTARY INFORMATION

**Figure S1.**
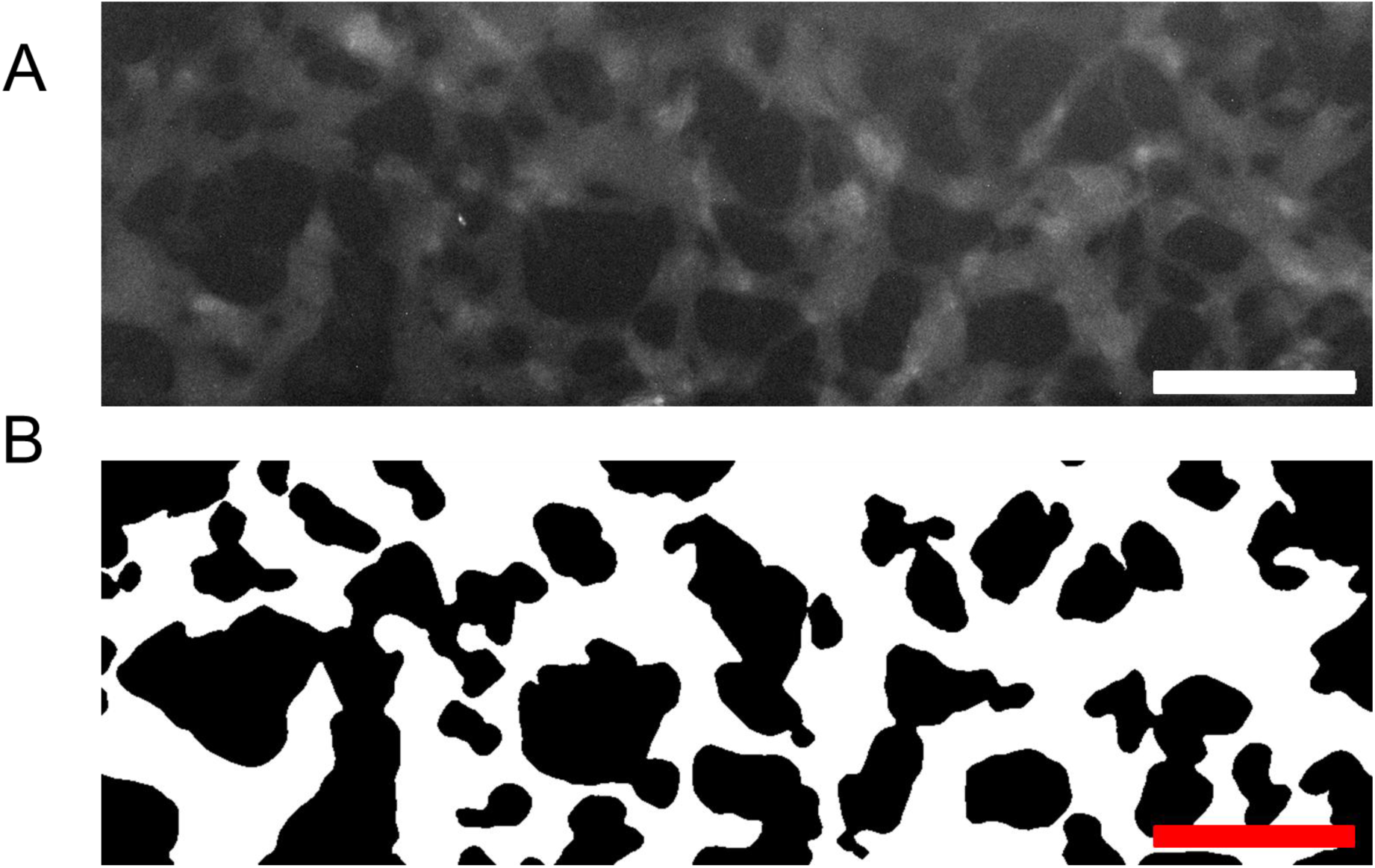
Measurement of vascular diameters. (A) Dextran channel of an image of a perfusable area of a microvasculature network on day 0. (B) Thresholded binary image of (A). Matlab Reaver was used to calculate the mean diameters. Scale bars: 500 µm.

**Fig. S2.**
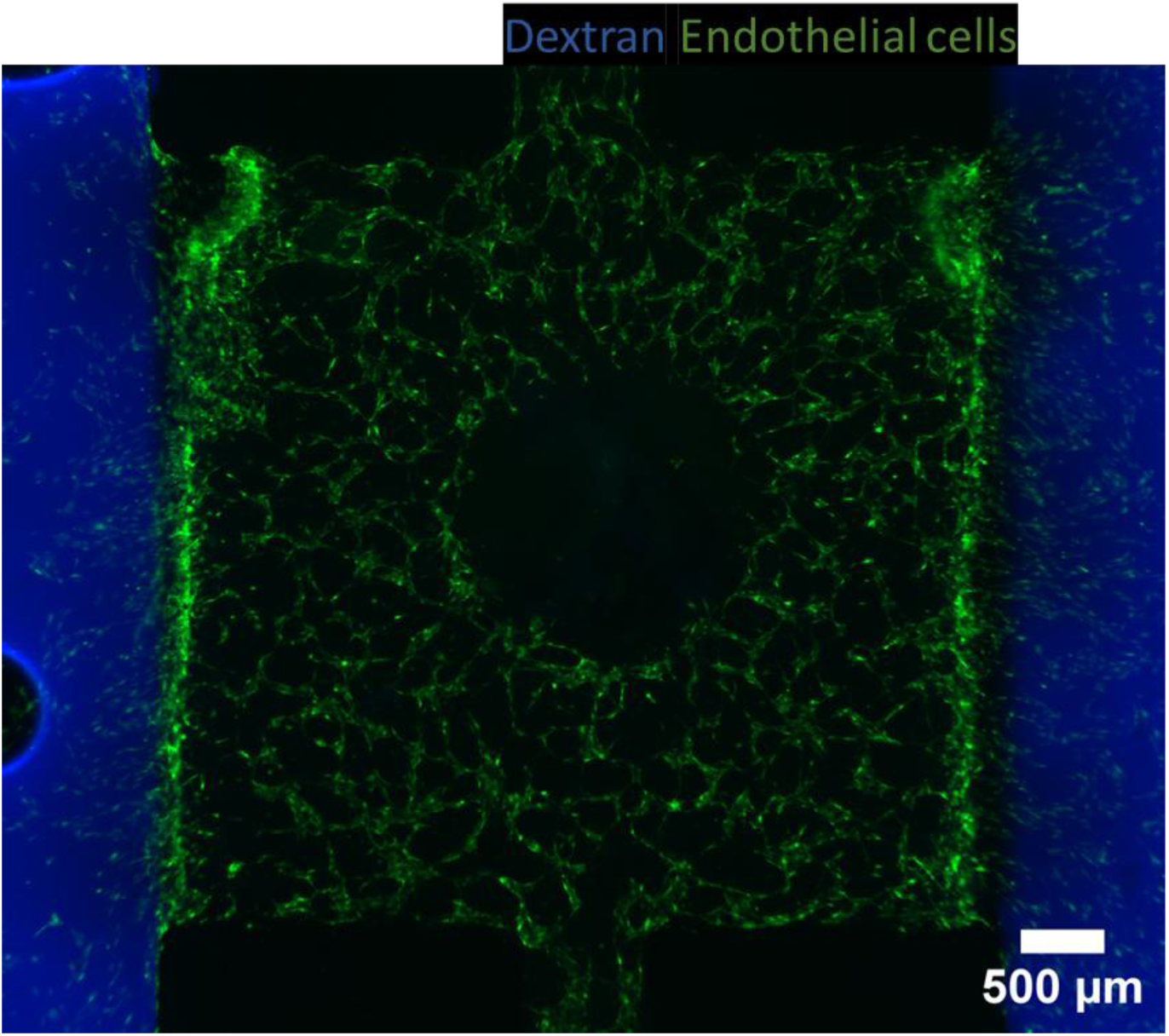
Devices created by co-culturing human umbilical vein endothelial cells (HUVECs) and mesenchymal stem cells (MSCs) The microvasculatures are less perfusable, partly due to the rapid growth of MSCs.

**Figure S3.**
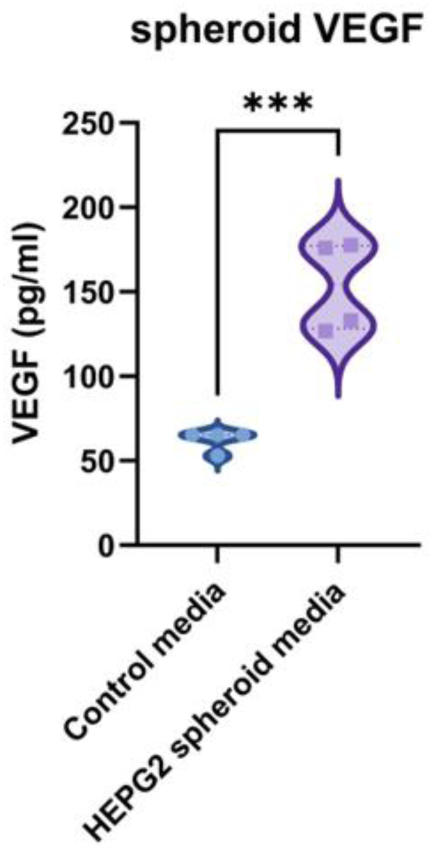
VEGF secretion by HepG2 in the culture media collected from well plates. The p-value was obtained from two-tailed Student’s t-tests.

**Fig. S4.**
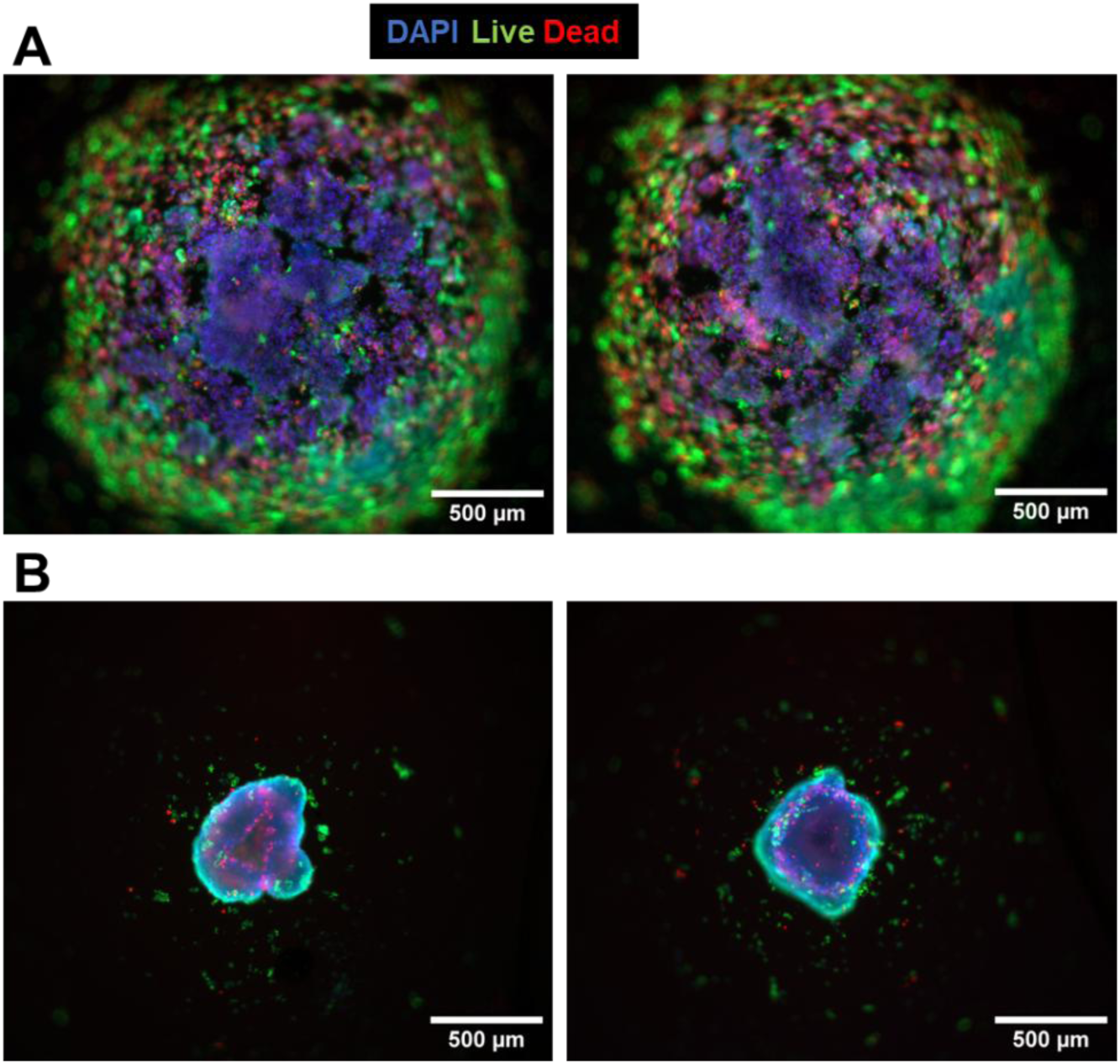
Spheroid morphology in the function of the number of seeded cells. A) Large spheroids with 1.3×10^4^ cells/well B) Small spheroids with 3333 cells/well.

## SUPPLEMENTARY VIDEO

**Video S1. Embolization of a vascularized tumor model using a bead-based embolic agent**

https://doi.org/10.6084/m9.figshare.28599905.v1

## Notes

### Competing Interest Statement

The authors have declared no competing interest.

